# Understanding *Lactobacillus paracasei* and *Streptococcus oralis* biofilm interactions through agent-based modeling

**DOI:** 10.1101/2021.04.29.441960

**Authors:** Linda Archambault, Sherli Koshy-Chenthittayil, Angela Thompson, Anna Dongari-Bagtzoglou, Reinhard Laubenbacher, Pedro Mendes

## Abstract

As common commensals residing on mucosal tissues, *Lactobacillus* species are known to encourage health, while recent findings highlight the pathogenic roles of *Streptococcus* species in these environments. In this study we used a combination of *in vivo* imaging experiments and computational modeling to explore biofilm interactions between *Streptococcus oralis*, an accessory pathogen in oral Candidiasis, and *Lactobacillus paracasei*, an organism with known probiotic properties. A computational agent-based model was created where the two species only interact by competing for space and nutrients. Quantification of bacterial growth in live biofilms indicated that *S. oralis* biomass and cell numbers were much lower than predicted by the model. Two subsequent models were then created to examine more complex interactions between these species, one where *L. paracasei* secretes a surfactant, and another where *L. paracasei* secretes an inhibitor of *S. oralis* growth. Further biofilm experiments support the hypothesis that *L. paracasei* may secrete an inhibitor of *S. oralis* growth, although they do not exclude that a surfactant could also be involved. This contribution shows how agent-based modeling and experiments can be used in synergy to address multiple species biofilm interactions, with important roles in mucosal health and disease.

**IMPORTANCE:** We previously discovered a role of the oral commensal *Streptococcus oralis* as an accessory pathogen. *S. oralis* increases the virulence of *Candida albicans* infections in murine oral candidiasis and epithelial cell models through mechanisms which promote the formation of tissue-damaging biofilms. *Lactobacillus* species have known inhibitory effects on biofilm formation of many microbes, including *Streptococcus* species. Agent-based modeling has great advantages as a means of exploring multifaceted relationships between organisms in complex environments such as biofilms. Here we used an iterative collaborative process between experimentation and modeling to reveal aspects of the mostly unexplored relationship between *S. oralis* and *L. paracasei* in biofilm growth. The inhibitory nature of *L. paracasei* on *S. oralis* in biofilms may be exploited as a means of preventing or alleviating mucosal fungal infections.

## INTRODUCTION

*Lactobacillus* and *Streptococcus* species are ubiquitous commensals found in the human oral cavity-but also the genitourinary and gastrointestinal tracts. Mitis group streptococci (MGS), primarily represented by *Streptococcus oralis, Streptococcus sanguinis, Streptococcus gordonii*, and *Streptococcus mitis*, are prominent among first colonizers of biofilms on mucosal and tooth surfaces (Diaz et al., 2006; Diaz et al, Mol Oral Microbiol, 2012). MGS were originally found to play a positive role, maintaining microbiome homeostasis in the oral cavity by antagonizing other microbes such as the cariogenic *S. mutans* (Manti et al., 2020; Thurnheer & Belibasakis, 2018). Although they are members of the healthy oral microbiota, MGS were more recently recognized for their role as accessory pathogens, enhancing the virulence of potentially harmful members of the microbiota, such as *Candida albicans* and *Porphyromonas gingivalis* (I. M.G. Cavalcanti, Del Bel Cury, Jenkinson, & Nobbs, 2017; Indira M.G. Cavalcanti, Nobbs, Ricomini-Filho, Jenkinson, & Del Bel Cury, 2016; Diaz et al., 2012; Jakubovics & Kolenbrander, 2010; Whitmore & Lamont, 2011; Xu, Jenkinson, & Dongari-Bagtzoglou, 2014). In addition, MGS can be pathogens in their own right: when they enter the bloodstream, they can cause endocarditis, bacteremia, and toxic shock (Shelburne et al., 2014; Douglas et al. 1993; Gassas et al. 2004).

Many *Lactobacillus* species have probiotic properties that promote gut, vaginal, and oral health (Mahasneh & Mahasneh, 2017; Santos et al., 2016; Reid et al., 2009; Guarino et al., 2015). *Lactobacilli* possess a diverse array of mechanisms implicated in the inhibition of vital processes in other bacteria; these include inhibition of growth through production of lactic acid and bacteriocins, and prevention of attachment to surfaces by competition, coaggregation and production of biosurfactants, which may also promote biofilm dispersion (Sharma & Singh Saharan, 2014; Tahmourespour & Kermanshahi, 2011; Kang et al., 2011; Haukioja, 2010; Satpute et al., 2016). In the oral environment, *Lactobacillus* species inhibit the growth and biofilm formation of *S. mutans* via multiple mechanisms. For example, secreted molecules found in supernatants of *Lactobacillus* cultures inhibit growth, adhesion and biofilm formation, and cell wall component lipoteichoic acid interferes with *S. mutans* sucrose metabolism, reducing the production of exopolysaccharide, an important component of biofilms (Ahn et al., 2018; Rossoni et al., 2018; Söderling et al., 2011). *Lactobacillus* spp also inhibit *Streptococcus pyogenes* hemolytic activity and adhesion to epithelial cells (Saroj, Maudsdotter, Tavares, & Jonsson, 2016). In addition to the great variety of antimicrobial effects attributed to different *Lactobacillus* spp, a considerable genetic and phenotypic diversity exists in oral streptococcal species, and even strains within the same species, which affects growth in different oral ecological niches and their role as pathobionts (Xu, Jenkinson, and Dongari-Bagtzoglou 2014). *L. paracasei* is known to produce molecules with bacteriocin and surfactant properties (Gudiña, Rocha, Teixeira, & Rodrigues, 2010; Ciandrini, E., Campana, R., & Baffone, W. (2017); Ciandrini et al., 2016) but interactions between *L. paracasei* and *S. oralis* in the biofilm have rarely been explored.

To fully understand the complex community interactions between species, mathematical modeling is a complement to an experimental approach (Song et al. 2014; Horn & Lackner, 2014). It helps consolidate data and after validation it can help in making predictions (Hellweger et al. 2016) and provide an integrative and quantitative understanding of the system studied. Agent-based models (ABM) are particularly suited to represent biofilms as they capture the activities of each individual cell (autonomous agent) in the community (Koshy-Chenthittayil et al., 2021). These models incorporate rules of growth, division, movement, and decay for each cell, and these rules can be deterministic or stochastic. The cells are embedded in a spatial environment with relevant physical constraints, such as diffusion of chemicals (Lardon et al., 2011; Bauer et al., 2017). The behavior of each individual agent and the environmental constraints contribute to the emergence of the total population behavior, i.e., the biofilm development and structure. Very few agent-based models have been constructed using input both from the literature and experiments [Sweeney et al. 2019; Martin et al., 2017]. Our model is another addition to this small group of ABMs built through crosstalk between experimentation and simulation.

This work aims to further our understanding of the interactions between *S. oralis* and *L. paracasei* during biofilm growth with a rarely used combination of agent-based modeling and experimentation. The agent-based model was a device to better understand the dual species biofilm growth characteristics. The growth parameters of the model were estimated using the experimentally determined behavior of single species biofilms and data from the literature. Live fluorescence imaging showed that the growth of *S. oralis* is inhibited in the dual biofilm with *L. paracasei*. We then constructed two models expressing two distinct hypotheses: non-competitive inhibition and surfactant production. The models were validated with further experiments to explore the nature of the interactions between *S. oralis* and *L. paracasei*.

## RESULTS

### Model calibration

We began by constructing a model where the only interactions between the two species were competition for space and for consumption of nutrients required for growth. Using the iDynoMiCS software (Lardon et al., 2011), we constructed an agent-based model with the two bacterial species competing for a carbon source and oxygen. The growth parameters of the two species with respect to nutrients were estimated using experimental data collected as follows. Each of the species was grown independently as a biofilm in different dilutions of the media for a period of 16 hours (figure S1). As expected, the biovolume of each species reduced according to the dilution factor. These data were then used to estimate the cell growth parameters for each species in the model with respect to consumption of the carbon source. The growth parameters of each species with respect to oxygen were taken from the literature (Van der Hoeven, 1989; Lardon et al., 2011). Simulations seeded with a similar relative density of cells of each species as used in experiments led to biofilms with properties depicted in Figure 1.

**FIG 1.**
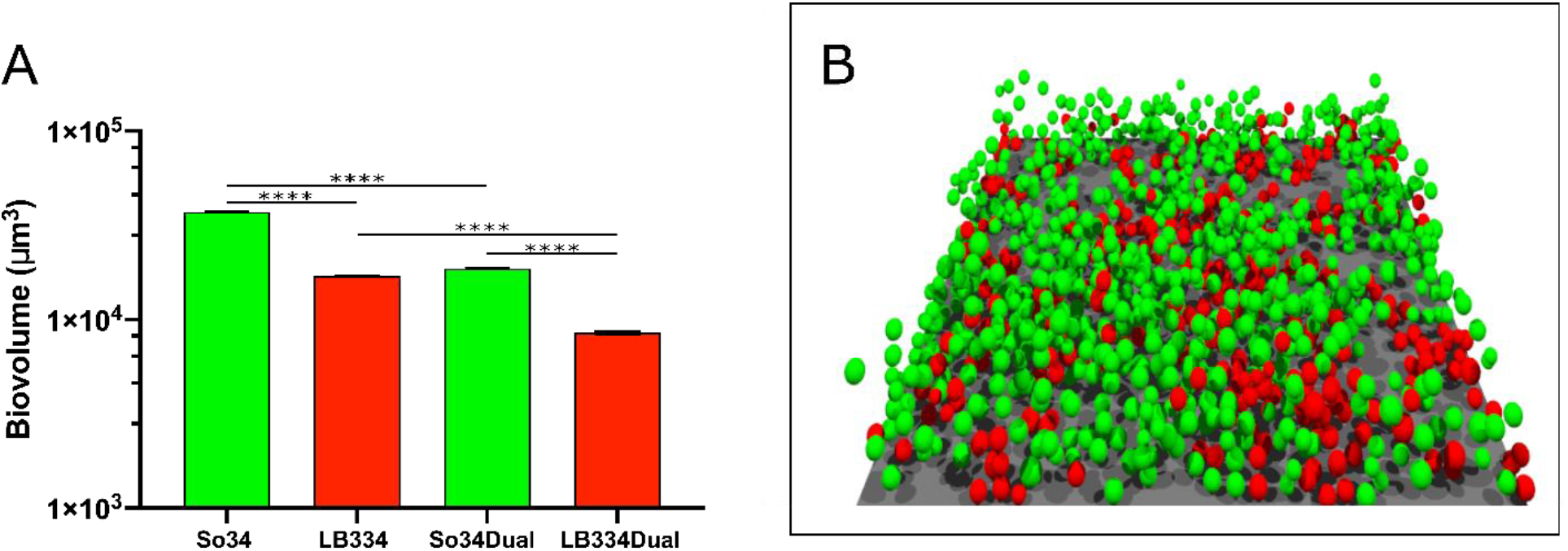
Simulation of biofilm growth in a purely competitive model. **A**- Biovolume for single and dual biofilms from simulations of 16-hour growth. Results are expressed as mean and standard deviation of 5 simulations with similar initial conditions. LB334- *Lactobacillus paracasei*, So34-*Streptococcus oralis*. **B**- Final structure of a 16-hour dual biofilm simulation. Red spheres- *L. paracasei* and green spheres- *S. oralis*; extracellular matrix filled the space between cells but is not represented in the image for clarity. The surface area of this image is 136 μm^2^.

### Model predictions

Agent-based models were set up to include each of the species in isolation, as well as a biofilm seeded with equal amounts of both species. These models were then run to examine how dual biofilms would behave under the hypothesis of simple competition for space and nutrients. As can be seen in Fig 1A, in single-species biofilms *S. oralis* is predicted to be a better biofilm former than *L. paracasei*, based on simulation biovolume estimates. As expected, there is a decrease in the biovolume of each species in the dual biofilm compared to the single biofilm, based on nutrient and oxygen competition. Interestingly, based on these parameters, the simulations predicted that in the dual biofilm the biovolume of *S. oralis* would be higher than that of *L. paracasei*.

### Live Biofilm Growth

We next tested single and dual biofilm growth of *S. oralis* and *L. paracasei* experimentally to validate the above model predictions. Each species attained a similar biovolume when grown alone for 16 hours, in contrast to the simulation, which predicted a lower biovolume for *L. paracasei*. In co-culture, *L. paracasei* growth was slightly lower than in single culture, but *S. oralis* biovolume was more significantly reduced (Figure 2A). This was in contrast to the simulation, which predicted a similar reduction in growth for both species. 3D projections of biovolume made from images of single and dual biofilms illustrate the altered growth pattern of *S. oralis* in biofilms with *L. paracasei* (Figure 2B-D). When growing alone, *S. oralis* biofilm takes the form of interconnecting mounds of cells and reaches thicknesses of 10-14 μm (Figure 2B). When growing with *L. paracasei, S. oralis* grows to a height of only 3-5 μm (Figure 2D, left image). *L. paracasei* maintains a similar growth pattern in single and dual-species biofilms (Figure 2C, D middle image). We used genus or strain specific qPCR of 16S rRNA to determine the number of each bacterial species present in the biofilms (Figure S2B). The qPCR method recapitulated what was observed for biovolumes: in dual-species biofilms, the number of S. *oralis* cells was significantly lower than in the biofilms containing *S. oralis* alone. Overall, these experiments indicated that actual biofilm growth did not match the predictions of the first iteration of our model in which competition for nutrients and space was the only interaction between the 2 species.

**FIG 2.**
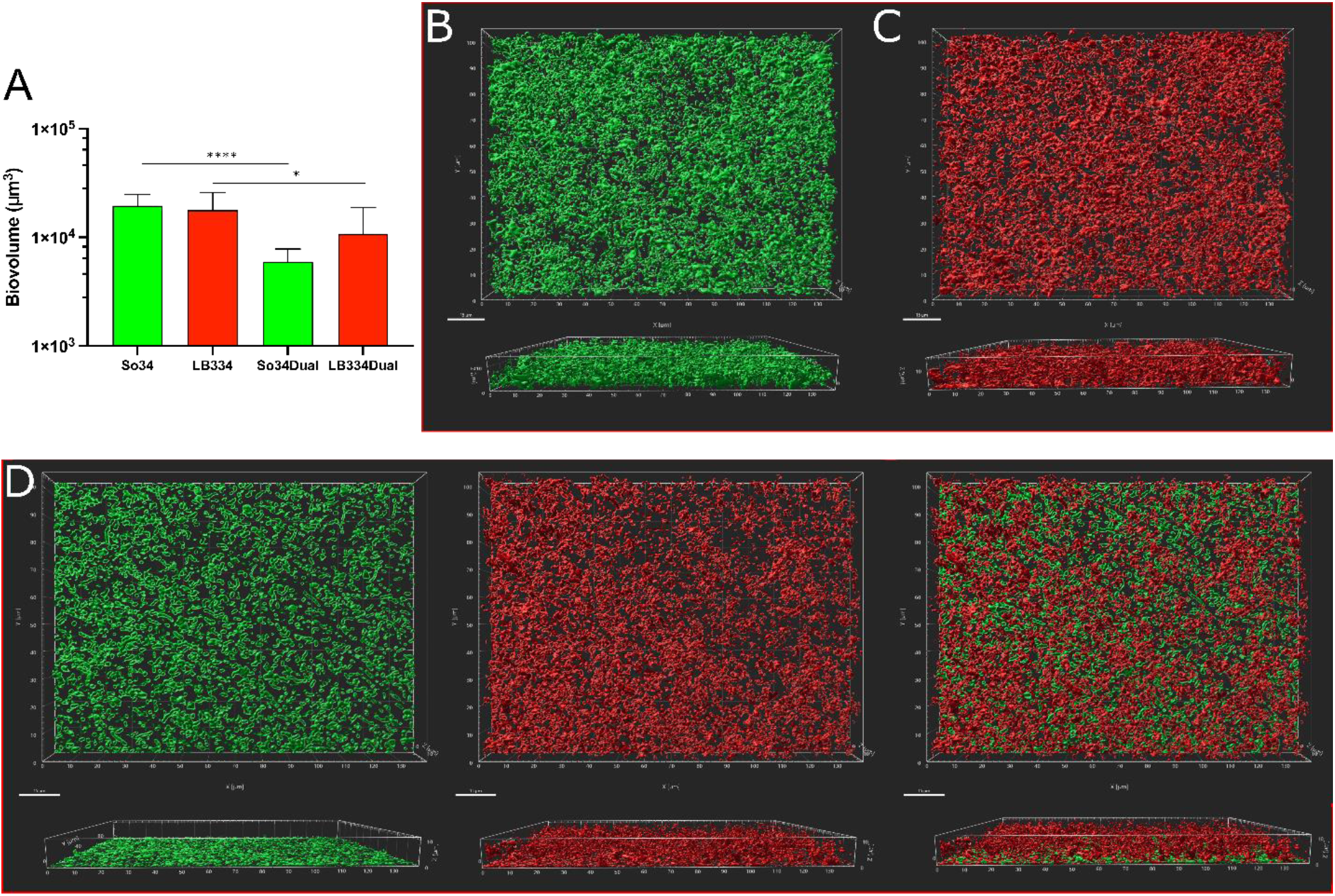
*In vitro* biofilm growth. **A**- Biovolume plot for 16-hour single and dual biofilm cultures. LB334- *Lactobacillus paracasei*, So34-*Streptococcus oralis*. Plotted is the average biovolume from 18 microscopic fields, imaged from 8 wells in 4 independent experiments. **B-D** Three-dimensional reconstructions of a 16 h *S.oralis* biofilm (B), a 16 h *L. paracasei* biofilm (C), and a 16 h dual *S. oralis-L. paracasei* biofilm *S. oralis*, left, *L. paracasei*, center, and merged 2-channel image, right (D). Scale bar is 15 *μm*

### Exploration of the possible interactions within the biofilm

Since we observed that there is a dramatic reduction in biovolume of *S. oralis* when it is grown with *L. paracasei* compared to when it is grown alone, we proposed two plausible explanations for these interactions:

a. *L. paracasei* product(s) may inhibit growth of *S. oralis* cells, or
b. *L. paracasei* may secrete a surfactant (Guidina et al., 2010; Ciandrini et al., 2016; Ciandrini, E., Campana, R., & Baffone, W., 2017) that causes both cell types to detach from the biofilm.

In the model, the growth inhibition mechanism was incorporated by including the production of a bacteriostatic product by *L. paracasei* and its effect on slowing the growth of *S. oralis*. To include a surfactant mechanism in the simulations, we used a modified version of iDynoMiCS (Sweeney et al., 2019) in a way that allowed the biofilm cells of both bacteria to disperse from the biofilm surface based on the local surfactant concentration (see Methods). We also ran a model which incorporated both mechanisms. The simulations were repeated 5 times each for the single and dual biofilm in all the three models.

In Fig 3, the growth inhibition model (3A), the surfactant model (3B) and their combination (3C) show that the biovolume of *S. oralis* is lower than *L. paracasei* in the dual biofilm suggesting that either or both mechanisms could be responsible for the decrease in *S. oralis* biovolume in biofilms with *L. paracasei*. In the combination model, we see that the *S. oralis* cells completely detach from the biofilm and thus its biovolume in the dual biofilm is zero (all *S. oralis* cells would end up detaching). Figures. 3D-F show the images of the simulated dual biofilm at 16 hours after inoculation. The images 3E and 3F depicting the simulation of the models with surfactant production, also show the floating planktonic cells that had detached from the biofilm.

**FIG 3.**
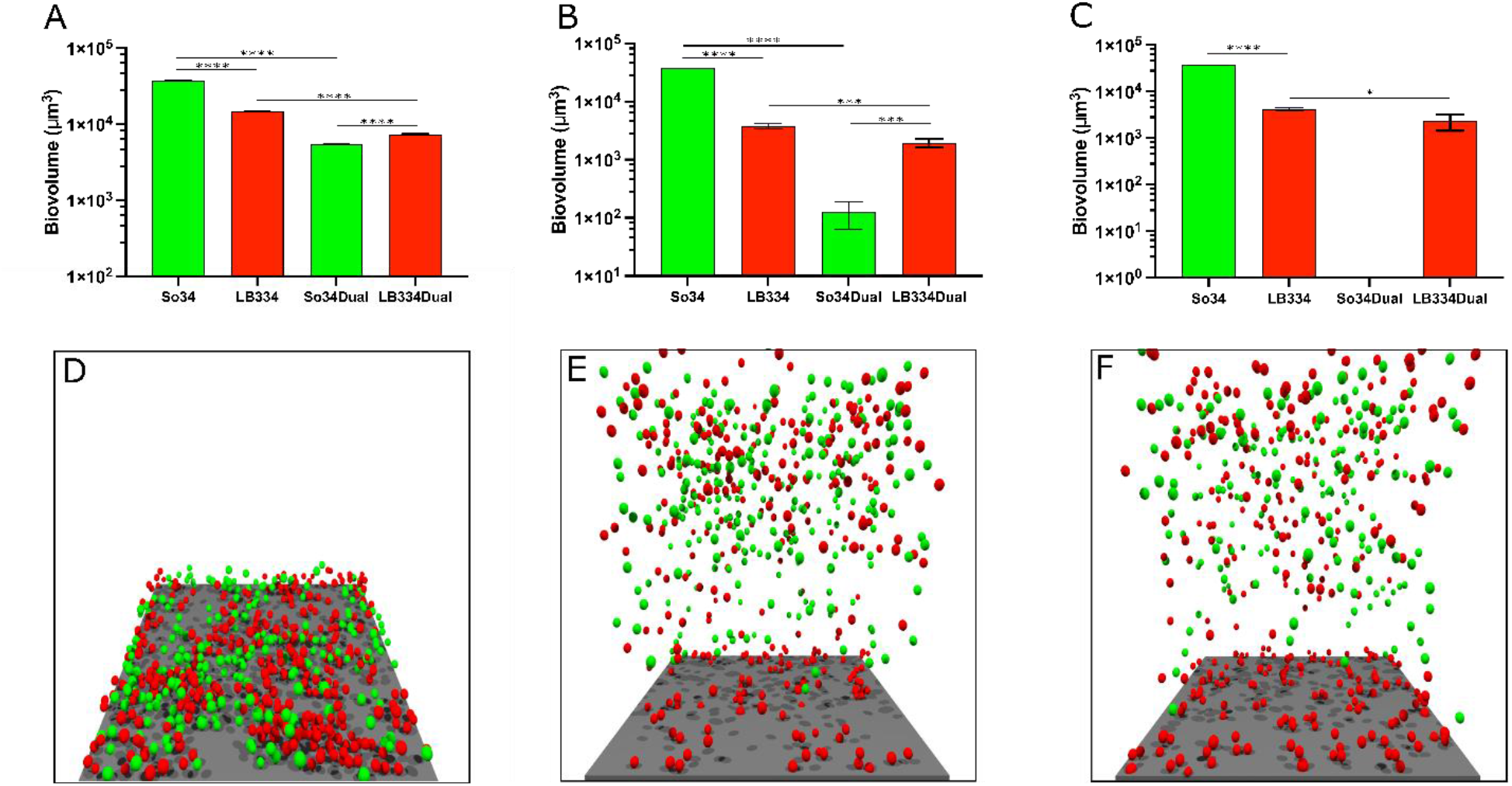
**A-** Biovolume plot for single and dual biofilms from the simulation of a 16-hour biofilm with growth inhibition. LB334- *Lactobacillus paracasei*, So34-*Streptococcus oralis*. Results are expressed as mean and standard deviation of 5 simulations with similar initial conditions. **B-** Biovolume plot for single and dual biofilms from the simulation of a 16-hour biofilm with surfactants. **C-** Biovolume plot for single and dual biofilms from the simulation of 16-hour biofilm with inhibition and surfactants. **D**- Final structure of a 16-hour dual inhibition model simulation. Red spheres- *L. paracasei* and green spheres- *S. oralis*; extracellular matrix filled the space between cells but is not represented in the image for clarity. The surface area of this image is 136 μm^2^. **E**- Final structure of a 16-hour dual surfactant model. The darker red and green spheres floating above the biofilm are the planktonic cells of *L. paracasei* and *S. oralis* respectively. **F-** Final structure of a 16-hour dual model with inhibition and surfactants.

Because the simulations of the two hypotheses resulted in somewhat similar outcomes (at least qualitatively), we are not able to eliminate either of them as possible explanations of how *S. oralis* is affected by *L. paracasei* in mixed biofilms. Thus, we looked for experimental evidence for the action of either an inhibitory substance, a surfactant, or perhaps both. Our culture wells contained both attached cells (in the biofilm) and planktonic cells (removed before imaging) and we reasoned that a surfactant or an inhibitory substance might alter the number of planktonic and biofilm cells in different ways. We expected that a toxin released into the surrounding media could reduce the number of cells in both the biofilm and the planktonic phase while a surfactant would tend to increase the proportion of planktonic bacteria without necessarily affecting the overall number of cells. To determine the number of cells in each phase, we enumerated each species using quantitative PCR with genus (for *Lactobacillus*)- or strain (for *Streptococcus*)-specific primers.

The total number of *S. oralis* cells in the biofilm, as counted by qPCR, was significantly reduced in the dual-species biofilms (Figure S1B). We determined that the number of planktonic *S. oralis* cells was also significantly reduced when *L. paracasei* was present (Figure 4A). *Lactobacillus* numbers were similar in single and dual-species biofilms by this quantification method, corresponding to our image analysis data (Figure 2A). Because the number of planktonic *S. oralis* cells was reduced along with the biofilm cells, we suspected a growth inhibitory toxin was present.

**FIG 4.**
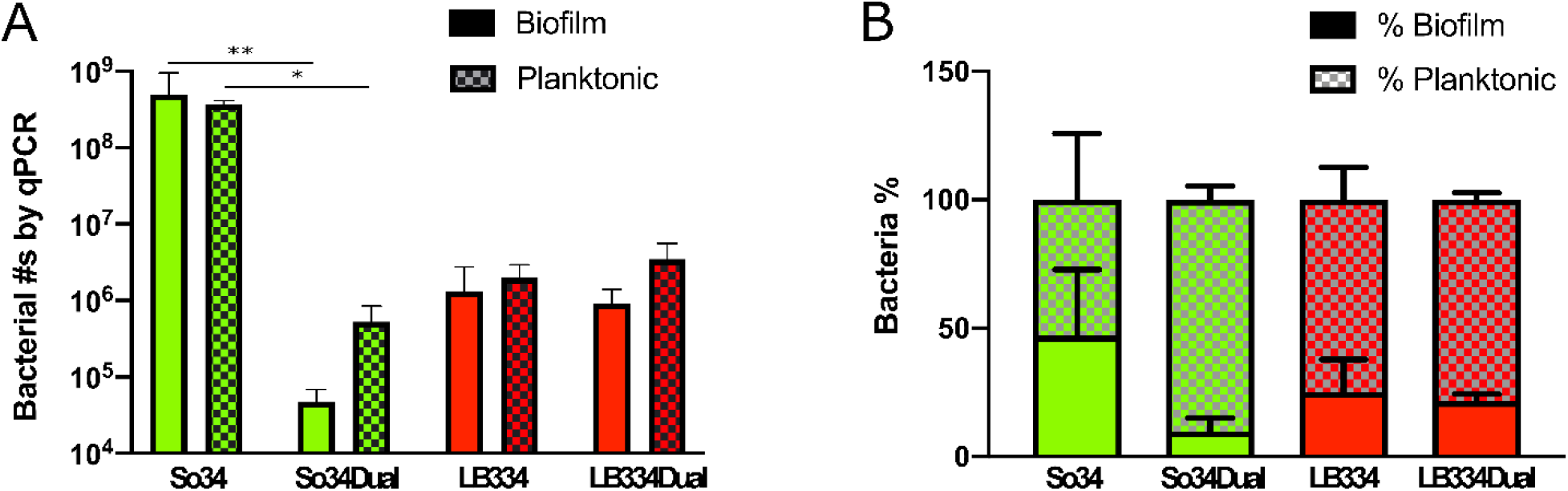
**A-** Plot of bacterial numbers in biofilm (solid bars) and planktonic state (hashed bars) for single species and dual species biofilms determined by qPCR. Average of bacterial numbers from 4 biofilms. LB334- *L. paracasei*, So34- *S. oralis* **B-** Data from 4A plotted as a percentage of the total bacteria per well.

In Figure 4B, the same data were plotted as a percentage of the total number of cells in each well. We reasoned that a surfactant might increase the percentage of cells that detached from the biofilm, regardless of the overall growth of the bacteria in the well. Interestingly, the percentage of *S. oralis* planktonic cells was higher in the mixed cultures than in the single cultures, while *L. paracasei* appeared to be unaffected, showing similar percentages of planktonic cells in single and dual cultures. Considering these results, the data suggest that the presence of surfactant activity is additionally plausible, indicating that more than one mechanism may be involved in the suppression of *S. oralis* biofilm growth in the presence of *L. paracasei*.

### Further Experimental Exploration of the System

To explore the potential *Lactobacillus* cytotoxic activity further, we treated pre-formed 16 h. *S. oralis* biofilms with concentrated, cell-free supernatants from *L. paracasei* biofilms for 22h. We then used a live/dead stain and fluorescence imaging to determine the viability of *S. oralis* in these biofilms. The proportion of live cells was not significantly different in biofilms treated with filtrate containing molecules smaller than 3 kDa, in unconcentrated media, or in concentrated media controls (Figure5A). Rhamnolipid surfactants from *P. aeruginosa* are known to have antimicrobial properties (Nitschke et al. 2005). Commercially prepared Rhamnolipids at 100 *μ*g/mL and 200 *μ*g/mL increased the proportion of nonviable cells in a dose-dependent (although not statistically significant) way (Figure 5A). Concentrated media from *L. paracasei* biofilms containing molecules larger than 3 kDa significantly reduced the proportion of viable cells in *S. oralis* biofilms (Figure 5A). These results suggest that one or more high molecular weight secreted products by *L. paracasei* could act as bacteriocins against *S. oralis* in mixed species biofilms. Figure 5B depicts live/dead staining of *S.oralis* cells in biofilms exposed to each condition.

**FIG 5.**
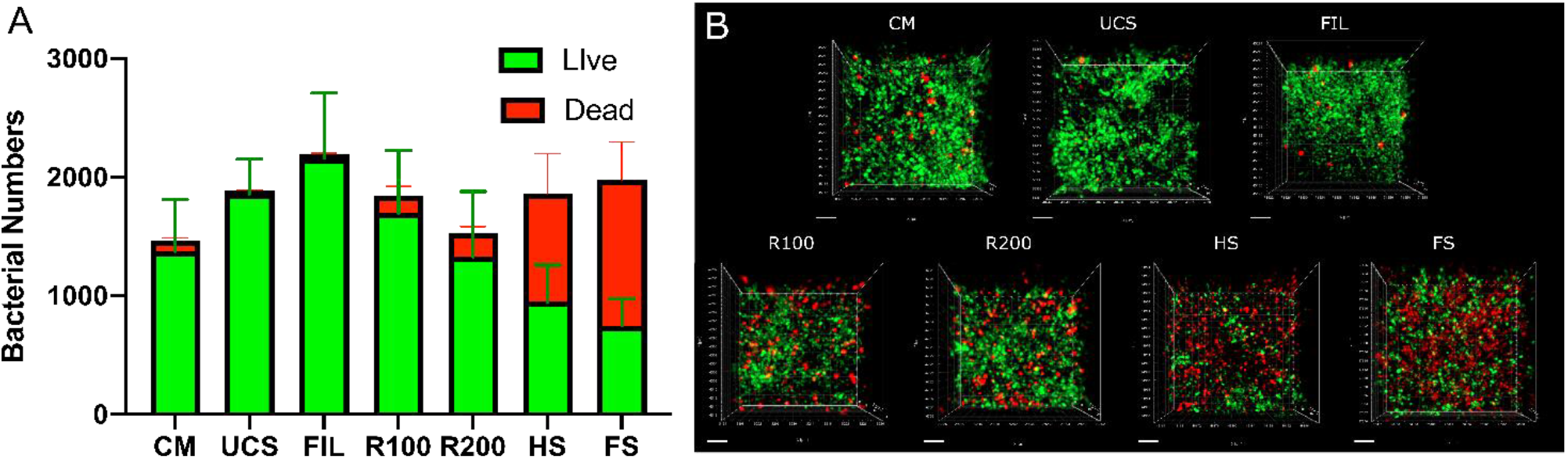
**A-** Number of live (green) and dead (red) *S. oralis* cells in biofilms exposed to CM-concentrated media control (n=15), UCS-unconcentrated supernatant (n=8), FIL-filtrate (n=13), R100- rhamnolipid positive control 100 *μ*g/mL (n=16), R200- 200 *μ*g/mL (n=8), HS-half-strength concentrated supernatant (n=17), FS-full-strength concentrated supernatant (n=15). Data is from two experiments and n denotes the number of images. **B-** 60x images (3-D) of live/dead staining of *S.oralis* cells in biofilms exposed to each condition. The scale bar is 5 *μm*.

## METHODS

### Bacterial Strains and Culture Methods

Fluorescent *S. oralis* 34 teal (Vickerman et al. 2015; Hongbin Xu et al. 2017) from glycerol stocks was grown overnight in brain-heart infusion (BHI) medium (Benton-Dickinson and Company, Sparks, MD, USA) supplemented with Erythromycin, 5 μg/mL, as needed, under aerobic static conditions at 37°C with 5% CO_2_. *L. paracasei* ATCC 334 (LB334) was similarly cultured in De Man, Rogosa and Sharpe (MRS) broth. Subcultures were grown from overnight cultures to an OD_600_ reading of 1.0 (1 x 10^8^ cells/mL). Prior to biofilm growth, LB334 were labeled with Cell-Tracker Red CMTPX dye (Invitrogen, Carlsbad, CA, USA) according to the manufacturer’s protocol. Briefly, cells were diluted 1:10 in PBS, centrifuged, resuspended at a density of 1 x 10^7^ to 1 x 10^8^ cells/mL in 18 μM dye in PBS, then incubated 45 min at 37°C. The labeled cells were centrifuged, washed once in PBS and resuspended in biofilm growth media. So34 cultures were diluted 1:10 in PBS. To satisfy the nutritional requirements of both bacterial species, biofilms were grown in an optimized complex medium containing 80% RPMI, 10% BHI and 10% Fetal Bovine Serum (Diaz et al, I&I 2012). 1 x 10^6^ bacteria were seeded into μ-Slide 8-well chambered coverslips (IBIDI GmbH, Gräfelfing, Germany), incubated at 37°C with 5% CO_2_ for 1 hour; the supernatant containing unattached bacteria was removed and replaced with fresh media, and incubation continued for 16 or 24 hours. Prior to imaging, media were removed and replaced with PBS.

### Microscopy and Image Analysis

Biofilms were imaged on a Zeiss Axio Observer inverted microscope with Apotome.2 (Carl Zeiss, Inc., Thornwood, NY, USA). Unless otherwise noted, images were made using a 63x oil immersion lens. Images were analyzed using Imaris software (Oxford Instruments plc, Tubney Woods, Abingdon, Oxon OX13 5QX, UK). Biovolume of biofilms was measured from 3D reconstructions using the “surfaces” protocol and live and dead cells were counted using the “spots” protocol in Imaris. Briefly, to measure biovolumes, we manually set a threshold intensity which excluded background fluorescence, then created “surfaces” which represent the volume occupied by each bacterial species. Objects less than ~1 um^3^ were excluded and the output volumes were added to obtain the total biovolume per image. To count individual live and dead cells from images, the diameter of spots (cells) was estimated to be 0.8 um, background subtraction was applied to the images and the total number of spots was recorded.

### qPCR Enumeration of Biofilms

Biofilms were grown as described above. Supernatants were removed at 1 h and 16 h, placed in sterile 2 mL tubes, centrifuged 5 minutes at 10,000 x g, and pellets were frozen at −80C. Biofilm starting cultures (1 h) and mature (16h) biofilms were frozen in their imaging well slides immediately after imaging. DNA extraction was carried out using the DNeasy Blood and Tissue Kit (Qiagen, Germantown, MD, USA) according to manufacturer’s instructions, including the suggested pretreatment with enzymatic lysis buffer for gram-positive bacteria. Genomic DNA was eluted in 100 uL nuclease free water. qPCR was carried out using S. oralis 34 strain-specific primers (*wefA-wefH*, Forward: 5’-CATCAAG AACTTCTCGGAGTTG-3’, Reverse: 5’-CCACAGCTCCAGAATA TTTAGC-3’)(H Xu et al. 2014) and All Lacto primers (Forward: 5’-TGGATGCCTTGGCACTAGGA-3’, Reverse: 5’-AAATCTCCGGATCAAAGCTTACTTAT-3’)(Haarman and Knol 2006).

### Preparation of Concentrated Supernatants

LB334 biofilms were grown in polystyrene 6-well plates for 24h under static conditions at 37°C with 5% CO_2_. Supernatants were centrifuged at 2,200 x g for 10 minutes, then filtered (0.2 ⍰m pore size). The remaining supernatant was concentrated by centrifugation in Centriprep centrifugal filters, 3,000 NMWL (Milipore) for 2 spins of 30 minutes, then 10 minutes at 3,000 x g. Filtered but un-concentrated supernatant, concentrated supernatant, and filtrate were frozen at −20C. Unconditioned media was also concentrated and used as a control. Supernatants were applied to 16h pre-grown So34 biofilms for 22h. The biofilms were then stained with a LIVE/DEAD BacLight Bacterial Viability Kit (Invitrogen) and imaged as described above.

### Data Analysis and Statistics

The biovolume and qPCR data were graphed, and statistical analyses were carried out in Prism9 (GraphPad Software, San Diego, CA, USA). Live biofilm data were analyzed by ordinary one-way ANOVA with Sidak’s multiple comparisons test.

### Agent-based model and simulations

All simulations shown were carried out with the iDynoMiCS software (Lardon et al., 2011) which can run agent-based simulations of biofilms including multiple species. The models in this paper include two bacterial species, *S. oralis* and *L. paracasei*, which are represented by two different types of agents in iDynoMiCS. These agents are governed by rules that represent cell growth, cell division, cell death (Figure 6), production of extracellular polymeric substances (EPS), cell movement through mechanical action (Figure 6) and cell detachment from the biofilm. The software also simulates medium nutrient diffusion and a liquid phase above the biofilm. Post-processing of simulation results was carried out with R (R Core Team, 2020) scripts and images were rendered with the PovRay (Persistence of Vision Pty. Ltd., 2004) software. All the relevant code can be obtained from https://github.com/skoshyc/StrepLactoBiofilmModeling.

**FIG 6.**
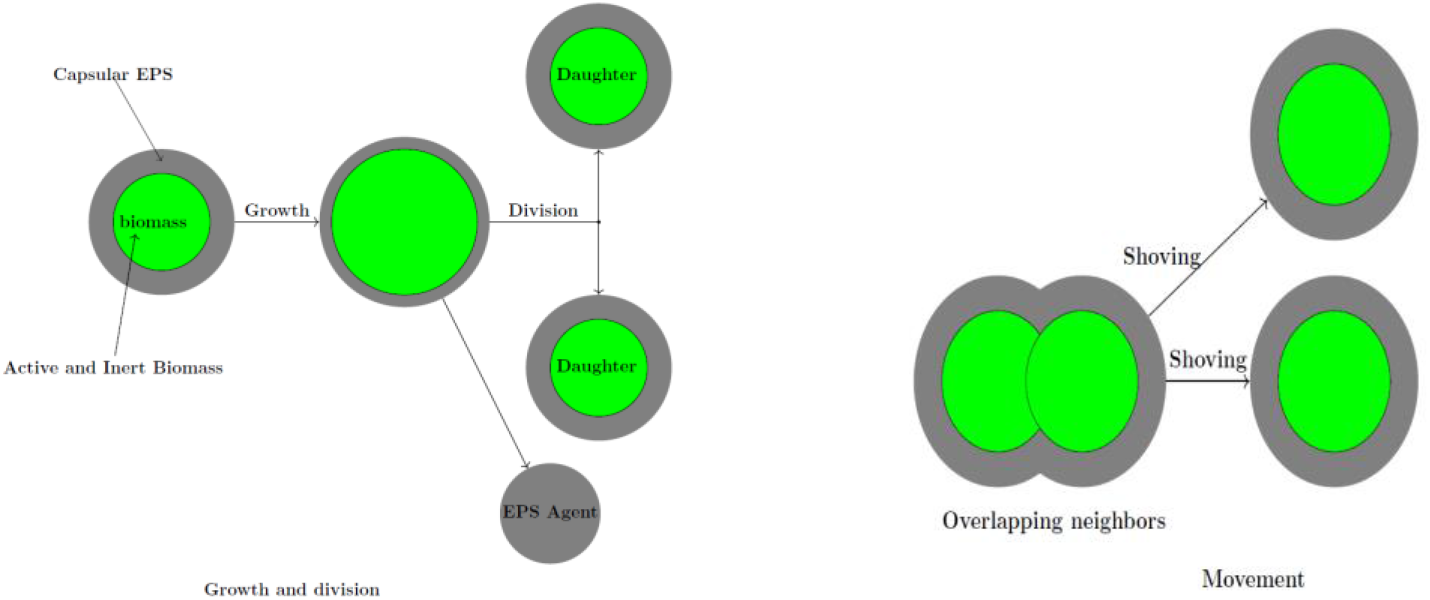
Growth, division, and shoving mechanism in the model based on iDynoMiCS (Lardon et al., 2011).

**FIG 7.**
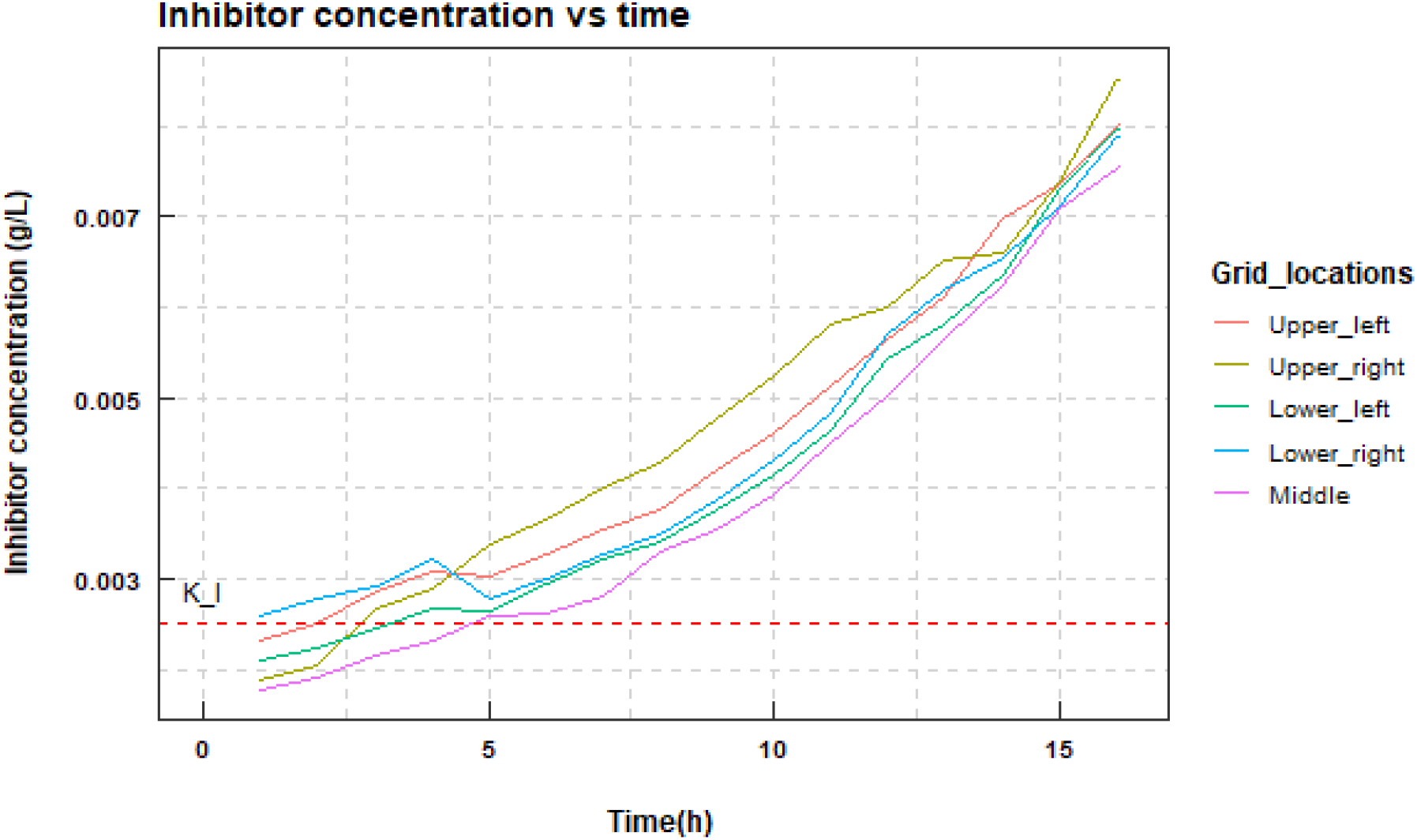
Inhibitor concentration in a dual biofilm at different time points in the simulation. Data were sampled at different locations of the biofilm, the four corners and the center of the surface area covered by the simulation. The red dashed line is the value of *K_I_* = 0.0025 which shows the inhibitory effect of the toxin on *S. oralis* is present from around 5 hours.

The models presented here are three-dimensional, with a computational grid of size 136 × 136 × 136 *μm*.

The growth rate for each cell size is given by:

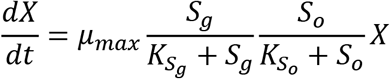

where X is the biomass of the cell. The two nutrients considered here are glucose and oxygen denoted by *S_g_* and *S_o_*, respectively. *μ_max_* is the maximum specific growth rate with unit *hr*^−1^ and the *K_S_* (unit is g/l) is the value of the substrate *S* when the specific growth rate is *μ_max_*/2.

### Estimation of growth parameters

We initially used growth parameters for each species obtained from the literature (Boudrant et al., 2005; Van der Hoeven, 1989; Rath et al.,2017; Lardon et al., 2011). The growth parameters specific to glucose were determined for planktonic growth of *Streptococcus gordonii* and *Lactobacillus casei* in a bioreactor. We then ran 16-hour simulations of single species biofilms for different glucose concentrations (2g/L, 1.5g/L, 1g/L, 0.5g/L). The initial value of 2g/L was set to match the sugar present in the biofilm media used in the experiments. On running simulations for different glucose concentrations, it was observed the biovolume of the species *S. oralis* and *L. paracasei* did not reduce with reduction in glucose, in contrast to what was observed in biofilm experiments. We assumed a biologically relevant range of the growth parameters related to glucose and applied a bisection method to estimate the desired parameter values that lead to reductions in biovolume upon reducing carbon source equivalent to experimental observations. The parameter values with respect to oxygen for each species were as in the literature (Van der Hoeven, 1989; Lardon et al., 2011). The final parameter values used in all the simulations are displayed in Table 1.

**Table 1 –.**
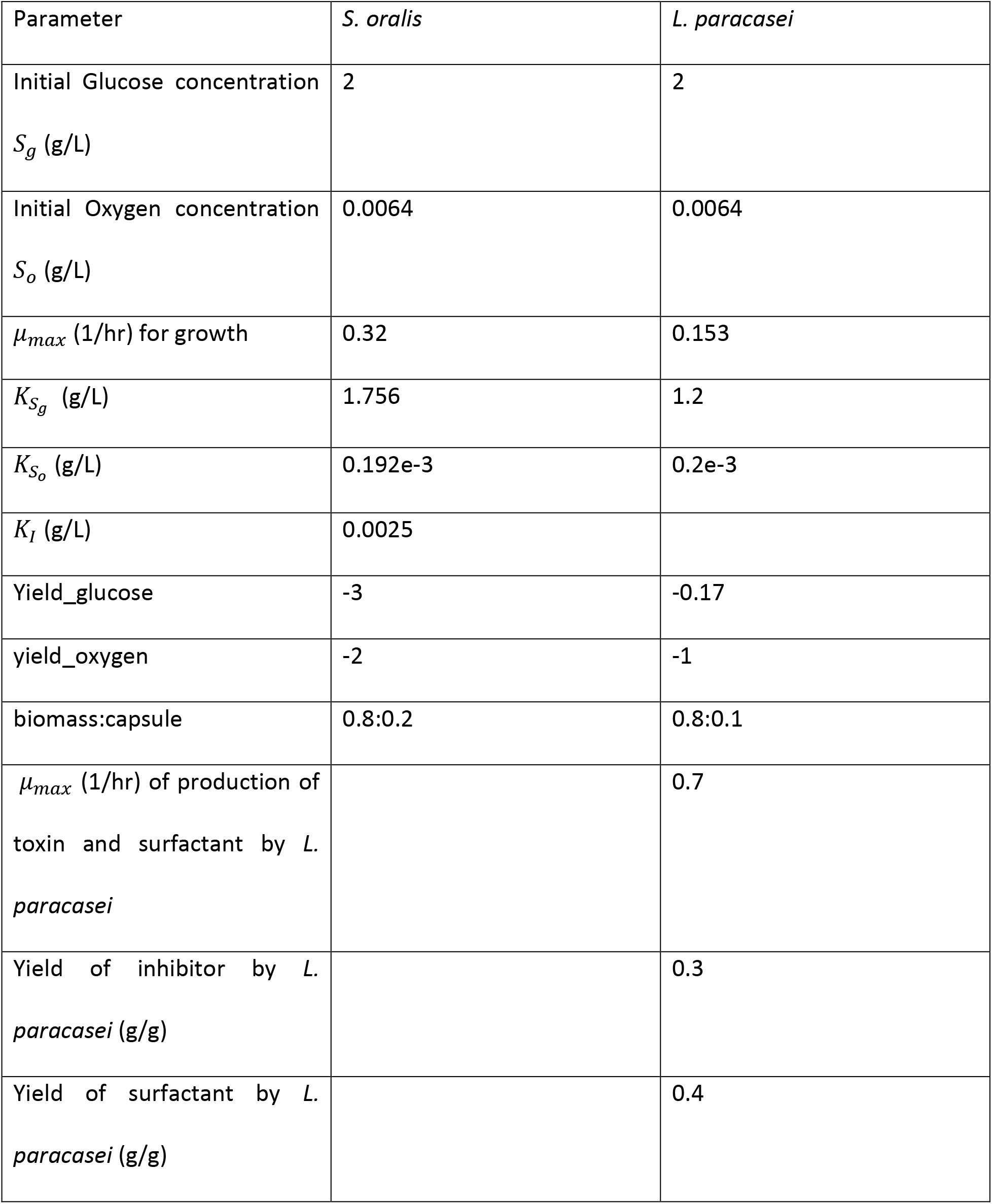
Parameter values used for cell growth of the two species in the biofilm simulations.

| Parameter | *S. oralis* | *L. paracasei* |
| --- | --- | --- |
| Initial Glucose concentration<br>$S_g$ (g/L) | 2 | 2 |
| Initial Oxygen concentration<br>$S_o$ (g/L) | 0.0064 | 0.0064 |
| $\mu_{max}$ (1/hr) for growth | 0.32 | 0.153 |
| $K_{S_g}$ (g/L) | 1.756 | 1.2 |
| $K_{S_o}$ (g/L) | 0.192e-3 | 0.2e-3 |
| $K_I$ (g/L) | 0.0025 | |
| Yield_glucose | -3 | -0.17 |
| yield_oxygen | -2 | -1 |
| biomass:capsule | 0.8:0.2 | 0.8:0.1 |
| $\mu_{max}$ (1/hr) of production of<br>toxin and surfactant by *L.*<br>*paracasei* | | 0.7 |
| Yield of inhibitor by *L.*<br>*paracasei* (g/g) |  | 0.3 |
| Yield of surfactant by *L.*<br>*paracasei* (g/g) |  | 0.4 |

All biofilm simulations were started with a seed of 176 cells. In the case of mixed species biofilm simulations, there were 88 cells of each type, such that the same total number of cells is kept constant. This matches the experiments which seeded the biofilms at 0.01 cells/μm^2^. The iDynoMiCS protocol files for the different simulations have been included as supplementary material.

### Non-Competitive inhibition

In this case the simulation includes secretion of by *L. paracasei* of an inhibitor to the growth of *S. oralis* through non-competitive kinetics. This required the following modification to the growth rate equation of *S. oralis*:

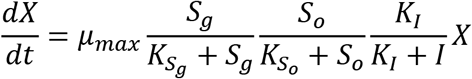

where *K_I_* is the concentration of the inhibitory substance *I* when the specific growth rate is *μ_max_*/2.

Since this is a hypothetical inhibitor, we cannot match its parameters to any real data. What is important is to have the *K_I_* for *S. oralis* to be around the concentration of that substance in the biofilm, so as to effectively cause an inhibition. We set *K_I_* arbitrarily to 0.0025 given the rate of production of the inhibitor by *L. paracasei* with a yield of 0.3. The inhibitor production is given as a separate first-order reaction with *μ_max_* = 0.7.

### Surfactant simulation

To simulate the action of a surfactant we utilized the software from (Sweeney et al., 2019) that is a modified version of iDynoMiCS 1.1 software. The original intent of their modification was to include secretion of a chemotaxis agent (Sweeney et al., 2019). In that modification the chemotaxis agent repels *Helicobacter pylori* cells which detach from the biofilm. For our purposes, the surfactant produced by *L. paracasei* plays a similar role to that repellant, however the effect is now on both cell types because the action of the surfactant is of a physico-chemical nature affecting both cell types (though not necessarily by the same degree, since that would be determined by their outer membrane composition). The surfactant is produced by *L. paracasei* cells and when its concentration becomes higher than a threshold, it causes cells of both species to detach from the biofilm and become planktonic.

Like in the inhibition model, we do not have real data on the tolerance levels of the bacteria to the surfactants. We assume that the *L. paracasei* require higher concentrations of surfactant before detaching from the biofilm (Gudiña et al. 2010). The tolerance threshold of surfactant for *L. paracasei* is 0.008 g/L and for *S. oralis* is 0.005 g/L. The thresholds were chosen based on the production of the surfactant by *L. paracasei* and were high enough so as to show a moderate effect (Fig. 8). The surfactant production is given as a separate first-order reaction with *μ_max_* = 0.7 with a yield of 0.4.

**FIG 8.**
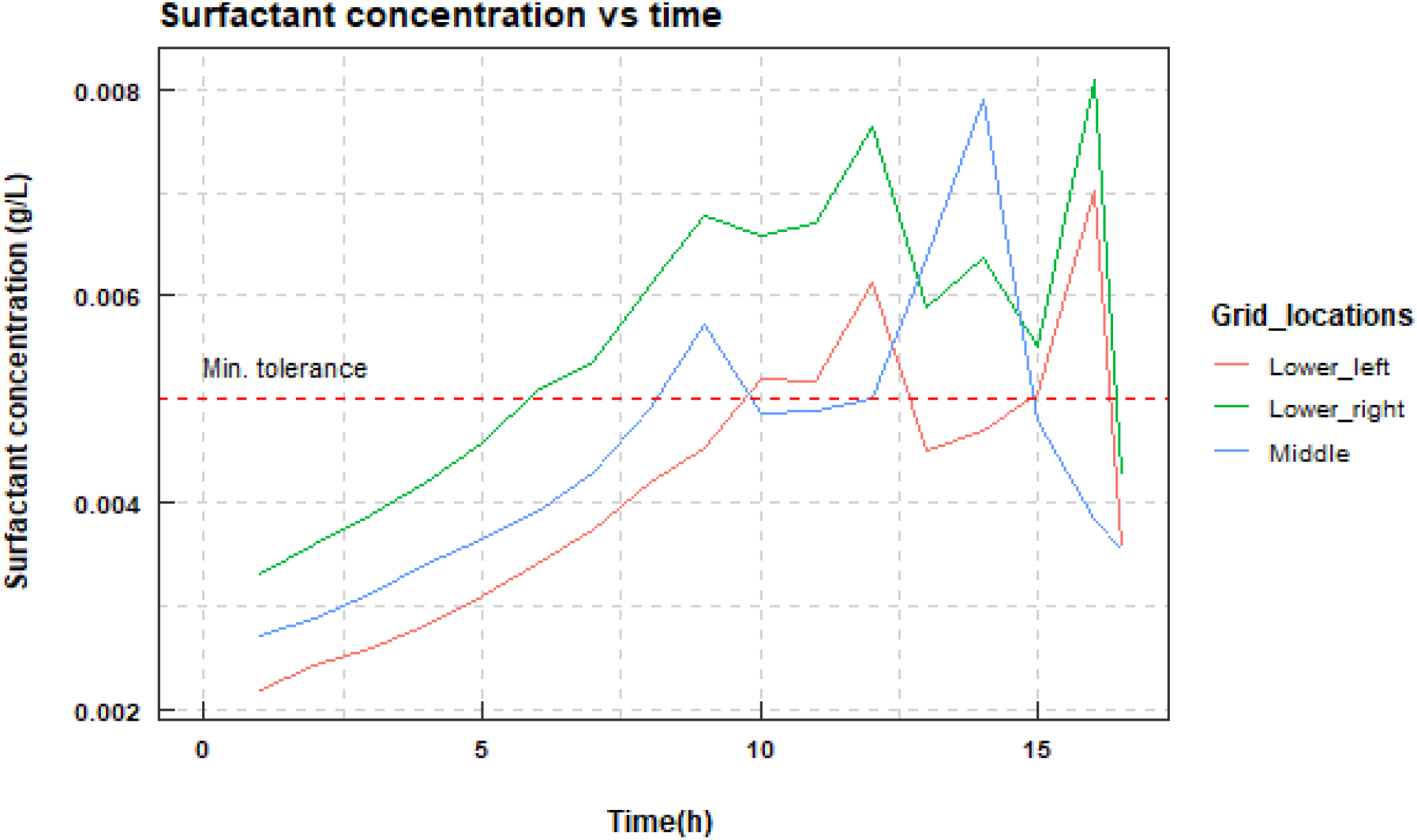
Surfactant concentration in a dual biofilm at different time points in the simulation. Data were sampled at different locations of the biofilm, the lower corners and the center of the surface area covered by the simulation. The red dashed line is the value of 0.005 which is the minimum of the tolerance thresholds for each of the species.

Unlike in the chemotaxis simulation of Sweeney et al. (2019), we assume that the planktonic cells do not rejoin the biofilm. To visualize this detachment in the simulation results, detached cells are depicted in a different intensity than the biofilm cells (Fig. 3E-F). The software includes random movement of planktonic cells in the liquid phase. The modified software that we used is available from https://github.com/alexwweston/iDynoMiCS.

We ran simulations of 16 hours of growth for both single and dual-species biofilms to see the effect of surfactants on the biofilm characteristics. We also ran simulations of the effect of the inhibitory substance and the surfactant together on the dual species biofilms. In all cases, statistics are provided of five simulations with the same input parameters, due to the stochastic nature of the simulations.

For each of the simulation models, the biovolume and species count was estimated using the R package iDynoR (Alden and Kreft, 2014). The average biovolume of the single and dual biofilms were compared using the Games–Howell test in the R package, ggstatsplot (Patil, 2018).

## DISCUSSION

We have used a combined modeling and experimental process to construct and refine a model of dual-species biofilm growth which has, in turn, informed further experimental exploration of interspecies interactions in biofilms. Our initial agent-based model that was limited to nutrient and space competition did not fully explain the interaction between *S. oralis* and *L. paracasei*. To account for the differences observed in the experimental results, we proposed two hypotheses for the inhibition of *S. oralis* biofilm growth: an inhibitory substance or a surfactant produced by *L. paracasei*. These hypotheses were translated into three different models and simulations were carried out with initial conditions similar to the experiments. In all three cases, the biovolume and cell numbers of *S. oralis* were reduced in the dual biofilm simulations compared to the single biofilm simulations, indicating that the experimental results could have been caused by either mechanism or both. Differences in *S. oralis* planktonic and biofilm growth in single and dual-species biofilms *in vitro* also supported both mechanisms. We further explored the interaction experimentally and found that concentrated supernatants from *L. paracasei* biofilm cultures contained a substance with mass larger than 3 kDa, which was toxic to cells in *S. oralis* biofilms. Prior to this study, *L. paracasei* and *S. oralis* interactions in biofilms had not been studied experimentally, and this is one of the few studies using a combination of agent-based models and experiments to study inter-species biofilm interactions.

Members of the viridans *Streptococcus* spp. have putative pathogenic roles, contrasted with the probiotic properties of *Lactobacillus* spp. and interactions between members of these genera have long been of interest to the medical community. For example, many *Lactobacillus* strains and their cell-free supernatants have antimicrobial activity against Streptococcal pathogens including *S. mutans* (Rossoni et al. 2018; Söderling, Marttinen, and Haukioja 2011; Wasfi et al. 2018; Ahn et al. 2018) and *S. pyogenes* (Saroj et al. 2016) (Meurman and Stamatova 2018). With a few notable exceptions (Ahn et al. 2018), the mechanisms of inhibition is not fully understood. Interactions between *Streptococcus* and *Lactobacillus* are often species specific and while most examples in the literature involve *Lactobacillus* spp. inhibition of *Streptococcus spp*., at least one example of the reverse has been found (Moon and Reinbold 1976).

In another area of intensive research, biosurfactants produced by *Lactobacillus* spp. have been explored as a means of inhibiting biofouling in commercial applications and in prevention of biofilms growing on hard surfaces in the oral environment (E. Ciandrini, Campana, & Baffone, 2017; Eleonora Ciandrini et al., 2016; Satpute et al., 2016). Surfactants are amphiphilic molecules that reduce surface tension and interfacial tension, thus interfering with adhesive interactions between microbes and the surfaces to which they attempt to attach. Many studies have found biosurfactant activity in supernatants of *Lactobacillus* culture, which are sometimes fractionated by molecular weight and identified by Nuclear Magnetic Resonance (NMR) and Fourier Transform Infrared Spectroscopy (FTIR) (Sharma and Singh Saharan 2014; Sharma et al. 2014; Shokouhfard et al. 2015; Gudiña et al., 2015). Molecules produced by *Lactobacilli* that have been implicated as biosurfactants include proteins, glycoproteins, glycolipids, phospholipids, lipopolysaccharides, lipopeptides, glycolipopeptides, and fatty acids (reviewed in: Satpute et al., 2016). In a few cases, purifications have been carried out to isolate the biosurfactant activity to a single substance (Vecino et al. 2013; Velraeds et al. 1996; Fouad, Khanaqa, and Ch 2010). Many of the isolated mixtures and substances have characteristics of both biosurfactant and bacteriocin, consistent with our findings with spent media in this work; for example, *Lactobacillus spp*. produce substances that have antimicrobial and antiadhesive activities (E. J. Gudiña, Rocha, Teixeira, & Rodrigues, 2010; Ciandrini, E., Campana, R., & Baffone, W. (2017); Ciandrini et al., 2016). Biosurfactant preparations from *L. paracasei ssp. paracasei* A20 completely inhibited the planktonic growth of several pathogenic and nonpathogenic bacterial species (including *S. oralis* J22) at a concentration of 25 mg/mL and showed dose-dependent anti-adhesive properties against many of the same species (Gudiña et al. 2010).

Our experimental results support the presence of an antimicrobial molecule in concentrated supernatants from *L. paracasei* biofilm cultures while enumeration of planktonic and biofilm-associated cells hints at a mechanism that affected adhesion or dispersal of *S. oralis*. Biosurfactants can have antimicrobial as well as antiadhesive properties, which could affect both the growth and adhesion of organisms in biofilms (Gomaa 2013; Satpute et al. 2016). It is also possible that our concentrated cell-free *Lactobacillus* spent media contains more than one active anti-streptococcal compound. Further fractionation and purification of *L. paracasei* supernatants could lead to identification of the substance or substances responsible for the effects we observed on *S. oralis* biofilms.

Our agent-based model based in iDynoMiCS (Lardon et al., 2011) is one of very few models which have been constructed with constant interaction and feedback with experimental work (Sweeney et al. 2019; Martin et al., 2017). The growth parameters in the model were based on the growth of single-species biofilms in different concentrations of the media, and prior published values. We investigated further interactions, namely non-competitive inhibition and surfactant effects, based on the observed interactions of *S. oralis* and *L. paracasei in vitro*. The surfactant model was constructed using a modified version of iDynoMiCS (Sweeney et al. 2019) which incorporated the transition of biofilm cells to planktonic cells based on the local surfactant concentration. A limitation of the model was that the area considered was much smaller than that used in experiments, due to limitations on computational resources. However, the ratio of initial number of seed cells to surface area was the same in the experiments and simulations. Effectively, the simulations represent a small section of the biofilms grown experimentally but have similar characteristics in terms of biofilm thickness.

This study contributes to an emerging picture of the role of Lactobacilli in controlling biofilm growth of other health-associated bacterial species such as *S. oralis* (Thurnheer and Belibasakis 2018). In certain host backgrounds this species synergizes with *C. albicans* to increase the virulence of the fungus (Diaz et al. 2012; Hongbin Xu et al. 2016; H Xu et al. 2014; Souza et al. 2020). *L. paracasei*, on the other hand, inhibits the transition of *C. albicans* yeast to hyphal form (Rossoni et al., 2018). Interkingdom interactions such as those between oral bacteria and the fungus *C. albicans* hold great interest due to medical treatments or immunocompromised states resulting in fungal-bacterial dysbiosis, a contributing factor in human disease (Abusleme et al., 2018). Our next goal is to increase the complexity of our experimental and mathematical models by incorporating *C. albicans* to study interactions in three species biofilms. An assumption made in this agent-based model is that all the cells (agents) are spherical in shape. One avenue of future research will be to incorporate the rod shape (for *Lactobacilli*) and filamentous shape (for fungi) into the software and observe the effect, if any, that these cell morphologies may have on the biofilm structure and dynamics. The insights gained through the iterative process of modeling and experimentation in our study of *S. oralis* and *L. paracasei* interactions during biofilm growth will guide us in exploring more complex, multi-species biofilms.

## ACKNOWLEDGEMENTS

We wish to thank Margaret Vickerman for providing the *S. oralis* 34 teal fluorescent strain. This work was supported by NIH grant RO1 DE013986 and RO1 GM127909. R.L. was also partially supported by NIH Grants 1R011AI135128-01 and 1U01EB024501-01, and NSF Grant CBET-1750183.

## SUPPLEMENTARY FIGURES

**FIG S1.**
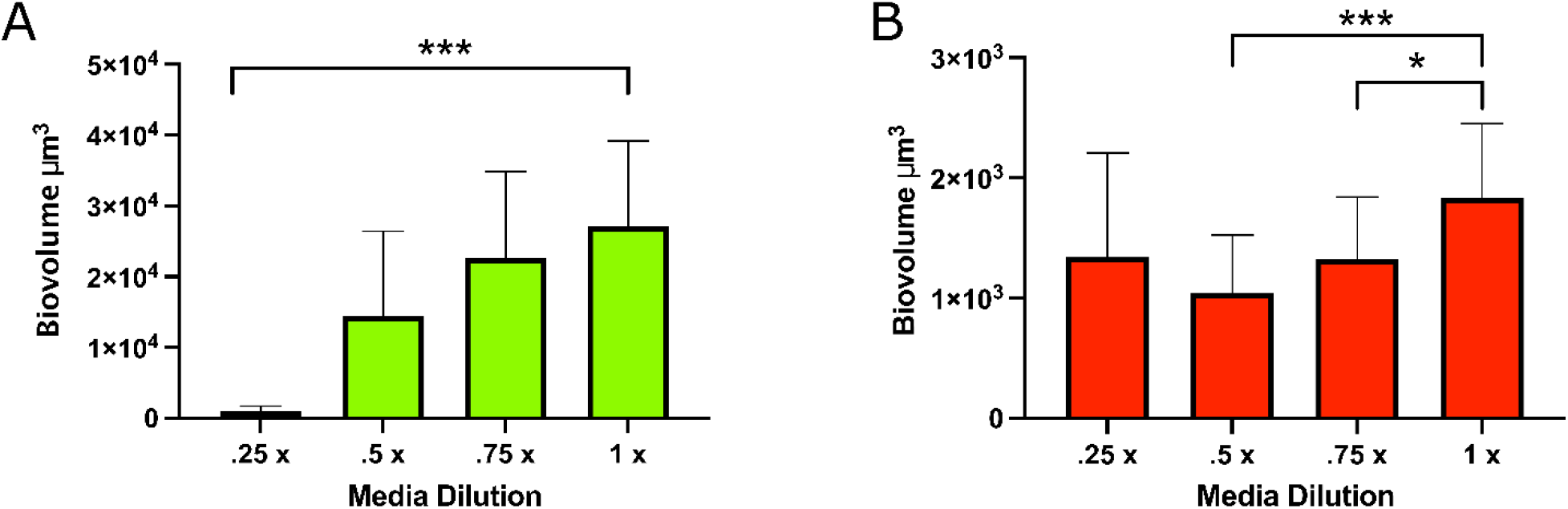
*In vitro* biofilm growth of *S. oralis* (A) and *L. paracasei* (B) in full strength biofilm media and media dilutions. Biovolume was measured in Imaris software, from images taken after 16h growth at 37°C, 5%CO_2_. *S. oralis:* 2 experiments, .25x media, n=5, .5x media, n=6, .75x media, n = 14, 1x media, n=16. *L. paracasei*, 3 experiments, n=19 for each media dilution.

**FIG S2.**
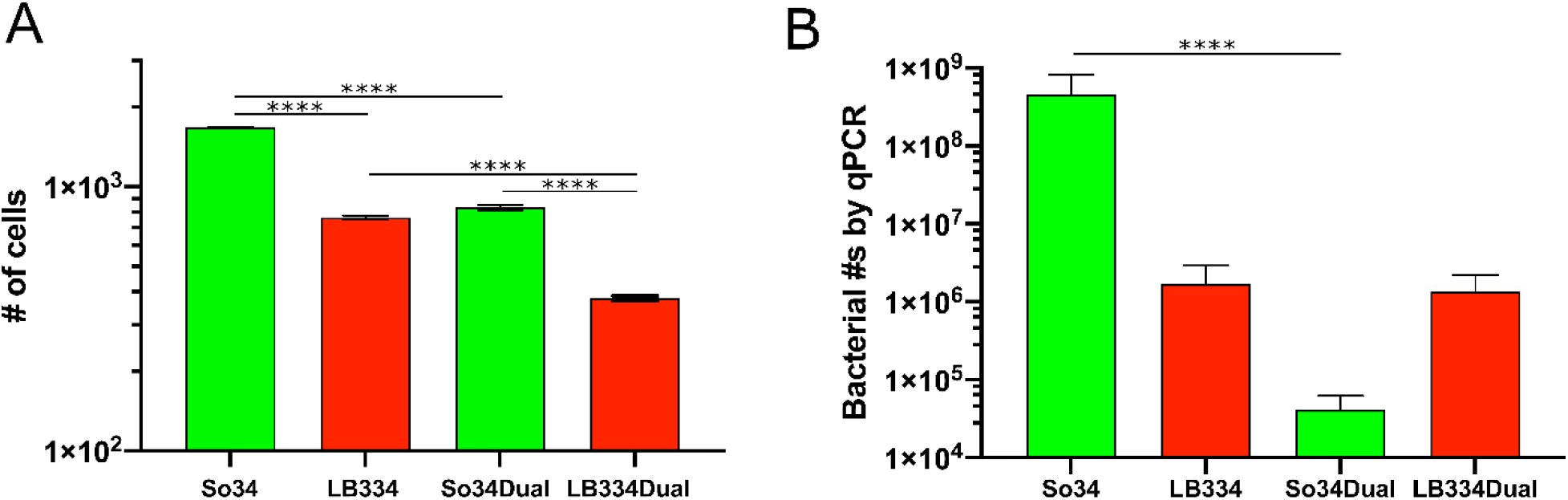
**A**- Number of bacterial cells (agents) for single and dual biofilms from simulations of 16-hour growth of a purely competitive model. Results are expressed as mean and standard deviation of 5 simulations with similar initial conditions. LB334- *Lactobacillus paracasei*, So34- *Streptococcus oralis*. **B-** Cell numbers in *in vitro* biofilms grown for 16h at 37°C, 5%CO_2_, as measured by qPCR targeting 16S rRNA gene for each species. Data are from 3 experiments, n=6 biofilms of each type: *S. oralis* alone, *L. paracasei* alone, and dual-species biofilm.

